# Dynamic task-belief is an integral part of decision-making

**DOI:** 10.1101/2021.04.05.438491

**Authors:** Cheng Xue, Lily E. Kramer, Marlene R. Cohen

## Abstract

Natural decisions involve two seemingly separable processes: inferring the relevant task (task-belief) and performing the believed-relevant task. The assumed separability has led to the traditional practice of studying task-switching and perceptual decision-making individually. Here, we used a novel paradigm to manipulate and measure macaque monkeys’ task-belief, and demonstrated inextricable neuronal links between flexible task-belief and perceptual decision-making. We showed that in animals, but not artificial networks that performed as well or better than the animals, stronger task-belief is associated with better perception. Correspondingly, recordings from neuronal populations in cortical areas 7a and V1 revealed that stronger task-belief is associated with better discriminability of the believed-relevant but not the believed-irrelevant feature. Perception also impacts belief updating: noise fluctuations in V1 help explain how task-belief is updated. Our results demonstrate that complex tasks and multi-area recordings can reveal fundamentally new principles of how biology affects behavior in health and disease.

## Introduction

Humans and animals make countless decisions every day that affect their well-being or even survival. In the laboratory, decision-making has typically been studied by observing behaviors and neuronal activity while subjects perform simple, well-understood sensory-motor integration tasks (Gold and Shadlen, 2007; Kable and Glimcher, 2009; Uchida, Kepecs and Mainen, 2006). But real-life decisions usually need to contend with a more important problem even before making perceptual judgements: inferring the relevant task to solve in a certain situation (i.e., task-belief). Task-beliefs allow decision-makers to deploy attention to a relevant subset of feature dimensions out of the huge amount of information in natural environments, and task-beliefs are flexibly adapted as the environment evolves (task-switching(Monsell, 2003)). Flexibly adapting task-belief is critical but difficult: the inability to appropriately respond to changing conditions is a debilitating symptom of disorders including autism, dementia, and substance abuse (Brady, Gray and Tolliver, 2011; Thapar *et al*., 2016; Dickstein *et al*., 2007).

Typically, task-belief is assumed to be a separate functional step that occurs before, and independent of, perceptual decision-making. In this view, subjective beliefs about the task (possibly involving parietal, prefrontal, and cingulate cortical areas (Stoet and Snyder, 2009; Buschman *et al*., 2012; Kamigaki, Fukushima and Miyashita, 2009; Sarafyazd and Jazayeri, 2019; Bartolo and Averbeck, 2020)) are used to identify task-relevant information. When that process is complete, the perceptual process (involving sensory areas such as visual cortex) makes a judgement about the believed relevant information (Purcell and Kiani, 2016a; Sarafyazd and Jazayeri, 2019; Mante *et al*., 2013) (Figure S1A, upper panel).

The goal of our study was to use behavioral, physiological, and modeling methods to test the view that flexibly inferring the correct task and perception are independent processes. To that end, we trained monkeys on a two-feature discrimination task that required them to report both their task-belief and their perceptual judgment on each trial. By decoding task-and perception-related signals from simultaneously recorded neuronal populations in parietal cortical area 7a and visual cortical area V1, we demonstrated that: 1) trial-by-trial fluctuations in task-belief strength correlate with perceptual performance and the fidelity of task-relevant information encoding in visual cortex; and 2) neuronal variability in visual cortex, even when it is unrelated to the visual input, plays an important role in updating task-belief (Figure S1A, lower panel). These results suggest an integrated neuronal system that serves as a common substrate for flexible task-belief and decision-making.

As neuroscience and cognitive science research start to shift focus from simplified functions in isolation towards more complex and natural behavior, our work has implications for methodology, basic science, and clinical translations at the time of this paradigm shift. Methodologically, our work showcases how dynamic cognitive states (e.g. task-belief) can be systematically manipulated and measured using large-scale neuronal recording from multiple brain areas during complex cognitive tasks. Scientifically, we demonstrated inextricable links between task-belief and perceptual decisions. These links would be impossible to see if each process were studied alone, highlighting that complex cognitive processes cannot be understood by investigating their components separately. Clinically, a wide variety of neuropsychiatric disorders lead to malfunctions in task-switching and decision-making. The neuronal link between the two processes suggests that potential treatments might do well to target neurotransmitters that mediate communication between brain areas.

## Results

### Perceptual decision-making during task-switching

Task-belief is an internal state that continually changes. Even with experimenters’ best attempts to keep task-belief constant (with fixed stimuli, explicit instructions, and task statistics), internal belief states still fluctuate substantially (Purcell and Kiani, 2016b; Ebitz, Tu and Hayden, 2020; Cohen and Maunsell, 2011a; Cohen and Maunsell, 2011b; Ashwood *et al*., 2022). These internal state fluctuations can impact visual cortical activity and perceptual decision-making (which is typically the only process that is measured in those studies; (Cohen and Maunsell, 2011a; Cohen and Maunsell, 2011b; Monsell, 2003). These results suggest that beliefs and perception interact in complex ways. The biggest barrier to understanding such interactions is that it is difficult to estimate task-belief during each decision because it is by definition internal and continually changing.

To address this challenge, we devised a novel two-feature discrimination task to assess perception and belief simultaneously and on every trial. We present to the animals two Gabor patches, one after the other. The second Gabor differs from the first in both of two features: spatial location and spatial frequency. The changes in both features are independently randomized. To obtain rewards, the animals were required to infer which of the two features as behaviorally relevant and to discriminate the direction of change in the relevant feature. Therefore, the task, requires two conceptually independent judgements to choose one choice target out of four (Figure 1A, upper panel). The animals were rewarded only when they chose the target corresponding to the correct direction of change in the relevant feature (Figure 1B, right panel). The relevant feature was not cued and switched with a low probability from one trial to the next (Figure 1A, lower panel).

**Figure 1.**
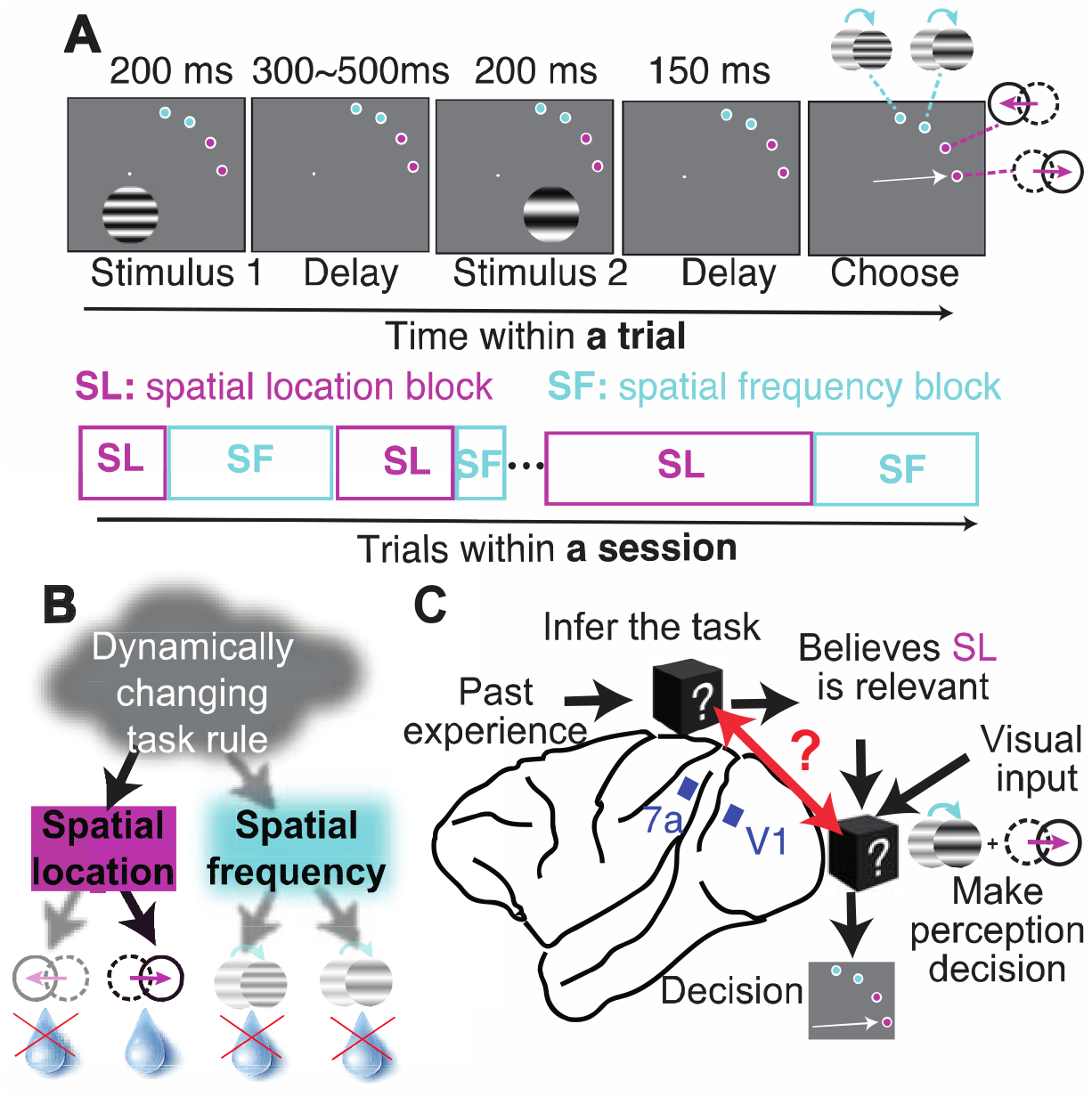
Behavioral paradigm and electrophysiological recording. (**A**) Schematic of the two-interval, two-feature discrimination task with stochastic task switching. On each trial, monkeys discriminate the difference in either spatial frequency or spatial location between two subsequent Gabor stimuli. The relevant feature is uncued and changes with 2.5% probability on each trial. The monkeys indicate their perceptual decision and the feature believed to be relevant by making a saccade to one of four choice targets. (**B**) and are rewarded for correctly reporting the sign of the change in the relevant feature. (**C**) Belief-based decisions could potentially be solved by independent hierarchical processes that compute belief and perception (black boxes). While the animals do the task, we simultaneously recorded population activity from one representative brain region related to each process (7a and V1 respectively, blue squares show approximate implant locations) to test the hypothesis that these processes are not separable (red arrow).

This design provides rich and easy measurements and manipulations of task-belief and perception through three key features: 1) on every trial, the animals make a perceptual discrimination, so we measure perceptual accuracy on every trial, regardless of whether the monkeys performed the correct task, 2) on every trial, the animals also explicitly indicate which feature is believed to be relevant, providing a behavioral ground truth about task-belief, and 3) by design, task-belief strength (the confidence that a chosen feature is behaviorally relevant) is overall stronger following a rewarded trial than following an unrewarded trial, providing a qualitative comparison with which to understand the implications of fluctuating belief state.

Another key aspect of our approach was to use neural population responses to decode continuous measurements of each stimulus feature and the monkeys’ task-belief on each trial. We therefore recorded from groups of neurons from which we could decode information about both visual features (in visual cortical area V1) and task-belief (in parietal area 7a (Kamigaki, Fukushima and Miyashita, 2009; Stoet and Snyder, 2004)) (Figure 1C). Together, these behavioral and physiological measurements provide a unique window into belief updating and perceptual decision-making on every trial.

The animals learned this task and performed it well. After training, the animals successfully discriminated the feature change they believed to be relevant and largely ignored the feature believed to be irrelevant (Figure 2A). They also effectively updated their belief according to the evolving task requirements, switching tasks only a couple of trials after the (uncued) task changes occurred (Figure 2B). The number of trials the animals took to notice task changes was close to optimal given their perceptual sensitivity (Figure 2C, see *Normative behavioral model* in STAR Methods for the optimal behavioral model).

**Figure 2.**
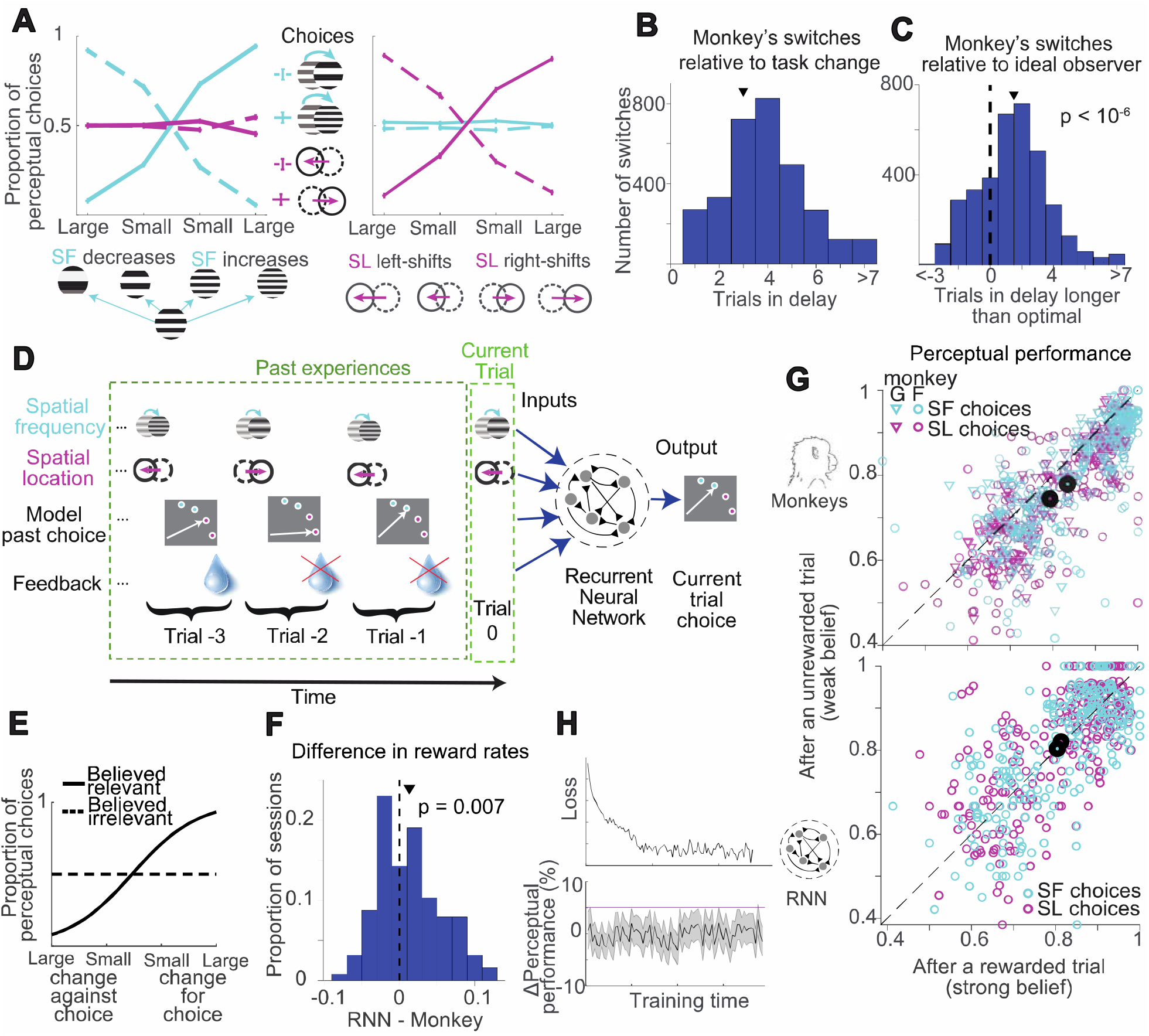
Trained behavior of animals and recurrent neural networks. (**A**) Psychometric curves showing the monkeys’ perceptual choice proportion as a function of spatial frequency (left panel) and spatial location (right panel) differences. The flat curves for the irrelevant feature show that animals successfully ignored irrelevant visual information. (**B**) Distribution of number of trials it took the monkeys to adapt to task changes across experimental sessions (mean 3.1 trials). (**C**) Distribution of number of trials the monkey took to adapt to the task change relative to an ideal observer model (see STAR Methods) (mean 1.5 trials). Positive values refer to occasions where monkeys were slower than the model; negative values indicate that the monkeys were accidentally faster. (**D**) Schematic of the inputs and outputs of the network. The network receives sequential inputs containing information about stimulus changes, past model choices, and choice feedback (reward history) over the course of the last several trials. Like the monkey, the network model is trained to infer the implicit task rule from recent history and make corresponding decisions. (**E**) Similar to the monkeys’ behavior in (A), the choices of a trained network are informed by the believed relevant feature information (solid line), while independent of the believed irrelevant feature information (dashed line). (**F**) Distribution of differences between reward rates between RNN and the monkeys across experiment session. Overall the RNN obtained more reward than the monkeys in the same experiment sessions. (**G**) Comparing perceptual performance following rewarded trials (abscissa) and unrewarded trials (ordinate) for monkeys (upper panel) and the recurrent network model (lower panel). Each point represents one stimulus condition of an experimental session, and we compute perceptual performance based on the subjectively chosen task, regardless of whether that task-belief was correct. Upper panel, the monkeys’ perceptual performance is better following rewarded trials than unrewarded trials. The distribution lies significantly below the unity line (p<10^−6^ for both monkeys and both features), showing lower perceptual performances following a non-rewarded trial than following a rewarded trial, with the same perceptual difficulty. Lower panel, the perceptual performance of the artificial network does not significantly depend on feedback history (p>0.05 for both features). (**H**) The difference between the monkey and network model is not explained by different extent of training. The upper panel shows that the loss function decreased during time, indicating the gradual learning process of the model on the task. In the lower panel, the black line indicates the network model’s difference between perceptual performance following rewarded and unrewarded trials at each point of training. The red line indicates the corresponding difference in the monkeys’ perceptual performance, which always lies outside the 90% confidence interval for the model (gray shading).

### Perception covaries with task-belief strength

Behaviorally, we found that dynamically changing task-beliefs affect the accuracy of perceptual decision-making. By design, the animals’ perceptual choices (as opposed to task choices) are informed by stimulus information within each trial and should ideally be independent from trial history. However, across experimental sessions (focusing on subsets of trials with the same stimulus conditions), the animals had better perceptual performance (i.e., perceptual accuracy of whichever task the animal chose to perform) after rewarded trials (which reinforced task-belief) than after unrewarded trials (which would weaken the monkeys’ task-belief; Figure 2G upper panel, Figure S2A).

Multiple lines of evidence suggest that the monkeys’ worse perceptual accuracy following a non-reward was due to a subsequent reduction in the animals’ certainty about their task belief. The perceptual worsening following non-reward is not explained by a potential tendency to make consecutive perceptual errors. Perceptual performance was still higher following perceptually-correct unrewarded trials (i.e. when subjects made correct perceptual decisions on the believed relevant feature, but the task-belief was wrong) than following task errors (Figure S2E). Further, the worsening of perceptual performance was even stronger following an easy perceptual decision without reward. Among trials following non-rewards, perceptual performance was worse when there was an obvious perceptual change in believed relevant feature, which comprised stronger evidence against the current task-belief, than when the feature change was ambiguous (i.e. weaker evidence against the task-belief; Figure S2F). Finally, the association between strong task-belief and improved perception persisted even when taking task-switch cost and task-set inertia (Alport, Styles and Hsieh, 1994) into account (Figure S2C-D). Correspondingly, the monkeys’ perceptual choices were more strongly related to V1 activity following rewarded than unrewarded trials (Figure S2B).

### Investigating the origin of the relationship between task-belief and perceptual accuracy

The observation that perceptual performance is related to task-belief can be explained by three broad, non-mutually exclusive hypotheses. First, it could represent a good strategy. This could occur if something about the structure of our task makes it more optimal to adopt a strategy that results in a relationship between perceptual performance and belief strength. Second, it could represent poor training or confusion, such as if the monkeys did not fully understand some aspect of the task. In this scenario, what we called weak task-belief would imply even more confusion, resulting in worse perceptual performance. Finally, this relationship could come from biology. The monkeys’ behavior could be constrained by a fundamental biological (potentially indirect) interaction between the neurons that encode task-belief strength and those that encode the visual features.

To investigate the plausibility of each of these hypotheses, we trained recurrent neural networks (RNNs) to perform the same behavioral task. Although RNNs learn very differently than real brains, they have the advantage that we could control the inputs, training history, and motivation (the model is optimized only to gain rewards, not for any other motivation, distraction, or effort that might influence human or monkey behavior). Like the monkeys, the RNNs had information about both the visual stimuli on the current trial and the recent trial history, and were required to infer the relationship between those inputs and the correct behavior (Figure 2D). We chose the amount of noise associated with the sensory input to the model so that the model and the monkeys have the same overall perceptual accuracy (Figure 2, A and E).

The behavior of the trained recurrent neural networks bears many similarities to the monkeys’ behavior. Like the monkeys, the network can distinguish relevant information from irrelevant information (Figure 2E) and can efficiently detect task changes and adjust behavior accordingly (Figure S3C). But unlike the monkeys, the network showed no relationship between perceptual performance and the reward history (Figure 2G).

These results are broadly inconsistent with the first two hypotheses. If the relationship between perception and belief were advantageous for receiving rewards, the enforced identical overall perceptual accuracy between monkey and model would mean that overall, the monkeys would receive more rewards than the model. We found the opposite: RNNs had higher reward rates than the monkeys (Figure 2F). If, in contrast, the relationship between perception and belief reflected the monkeys’ incomplete understanding of the task or poor training, we would expect the model to show this relationship early in training. We found the opposite: the network model never displayed a significant relationship between perceptual performance and past feedback during any stage of training (Figure 2H). These results demonstrate that better perceptual performance after rewarded than after unrewarded trials is neither a necessary aspect of performing this task, nor is it a necessary intermediate learning stage. Instead, they support the third hypothesis, suggesting that the difference between monkeys and models reflects the biological realities of how brains calculate task-belief and perception.

We therefore investigated potential neuronal interactions between task-belief strength and the representation of relevant and irrelevant features in visual cortex in V1. To test this hypothesis on a trial-by-trial basis, we looked for the neuronal correlate of task-belief in the activity of populations of neurons in area 7a of parietal cortex, which encodes task-related information (Stoet and Snyder, 2004; Kamigaki, Fukushima and Miyashita, 2009) and may mediate effects of trial history on perceptual choices (Akrami *et al*., 2018).

We reasoned that the worse perceptual performance following non-rewards might imply a moment-by-moment relationship between the monkeys’ internal belief state and the way the relevant and/or irrelevant visual feature is encoded in the brain. We therefore investigated how belief estimates in 7a relate to how visual information is encoded in V1 on a trial-by-trial basis. We used area 7a to decode our estimate of the animals’ belief state because, using a linear decoder, we were able to predict the animals’ choice of tasks significantly better from area 7a than from area V1 (Figure S1B lower panel, S1C).

We decoded a continuous measure of the animals’ task-belief from the population of neurons in area 7a on each trial (Figure 3A), and these estimates were related to behavior in reassuring ways. Consistent with the idea that rewards lead to stronger beliefs, the animals’ choice of which task to perform was better classifiable from area 7a after rewarded than an unrewarded trials (Figure 3B). Decisions to switch tasks were associated with a dynamic change in decoded task-belief away from the old task and toward the new task (Figure 3C).

**Figure 3.**
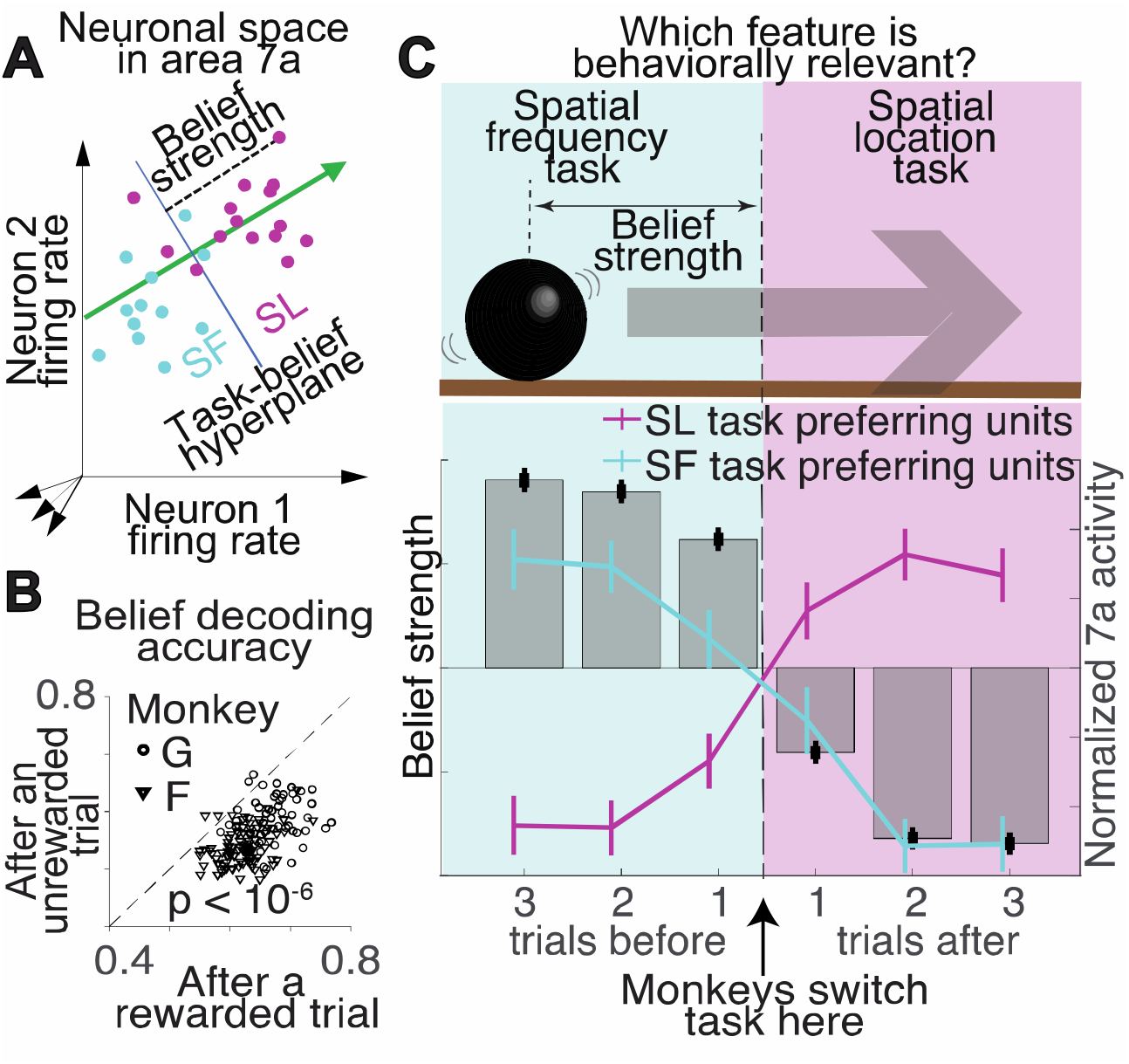
Neuronal measures of belief strength. (**A**) In a high dimensional neuronal space expanded by the activity of 7a units during the delay period, we find the best hyperplane to discriminate the task the animal performed on the trial. We define our single-trial neuronal measure of belief strength as the Euclidean distance from 7a population activity on each trial to the hyperplane. (**B**) Task-belief decoded from a neuronal population. Task-belief can be better classified by a linear hyper plane from area 7a population activity following rewarded than unrewarded trials. The abscissa and ordinate of each point show the performance of a linear classifier following rewarded and unrewarded trials respectively, with leave-one-out cross-validation. Each point represents one experimental session. (**C**) Belief strength is schematized as the distance from a rolling ball to a boundary. for trials leading up to the animals’ decision to switch tasks, the average belief strength decreased monotonically, changed sign right at the point the monkey decided to switch tasks and recovered as the new task-belief was reinforced (histograms in bottom panel). Normalized activity of task-selective 7a units tracked the same dynamics as decoded belief around task switches (lines in bottom panel). Error bars indicate standard errors.

We estimated the encoding of each visual feature on each trial using V1 population responses (Figure 4A), and these measurements were also related to behavior in intuitive ways. As expected, when comparing trials using the same stimuli (i.e. same difficulty), trials with larger relevant feature discriminability yield better perceptual performance (Figure S1E-F).

**Figure 4.**
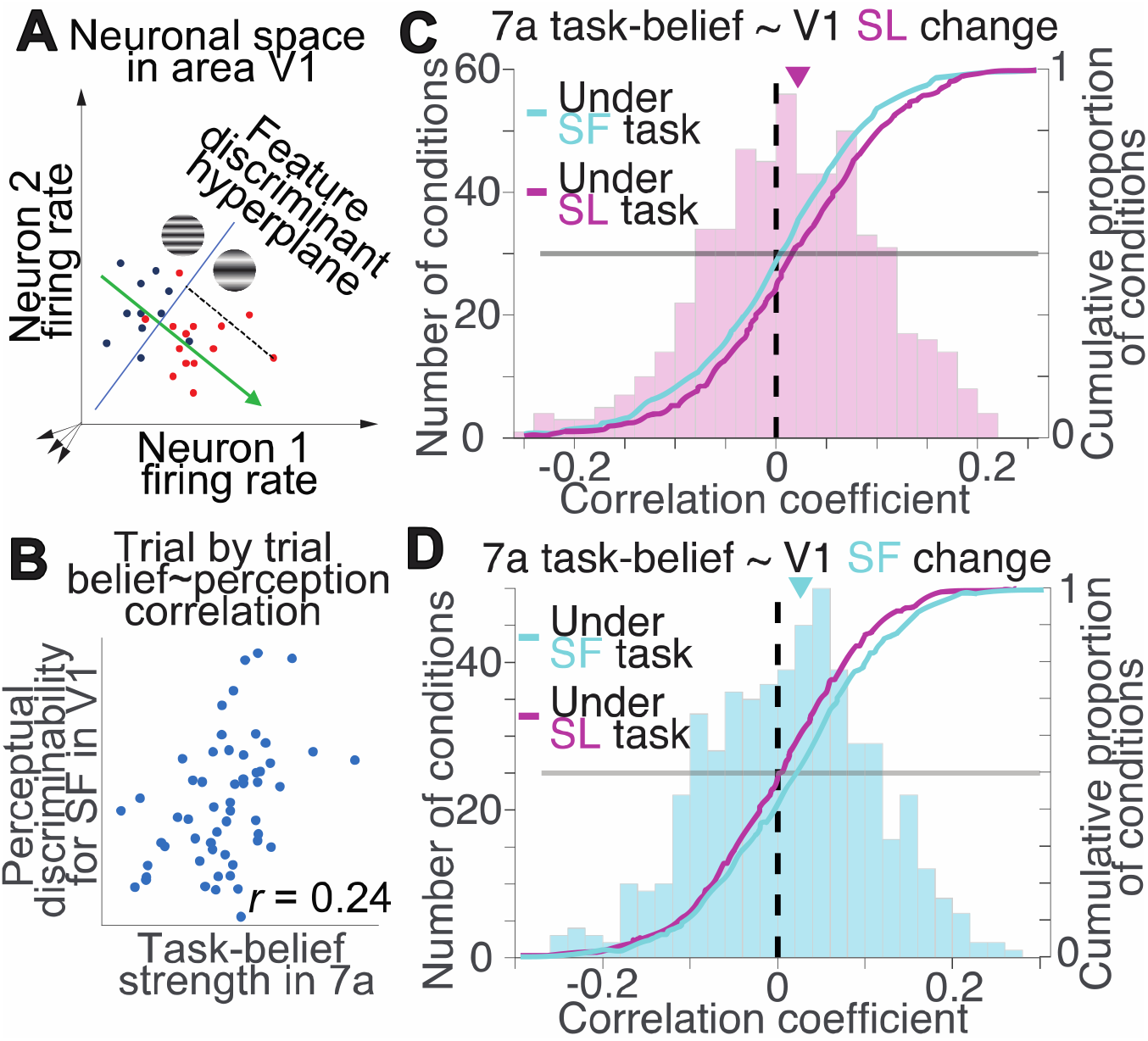
Belief and perception are linked on a trial-by-trial basis. (**A**) Using a procedure similar to that described in Figure 3A, we define the perceptual discriminability of each stimulus feature change on each trial as the Euclidean distance from V1 population activity to the hyperplane that best classifies the stimulus change of that feature (e.g., higher vs. lower spatial frequency). (**B**) Trial-by-trial comparison between belief strength (abscissae, decoded from 7a) and perceptual discriminability (ordinates, decoded from V1) for an example stimulus/task condition. If belief decisions and perceptual decisions are implemented by separate functional circuits of the brain, then internal fluctuations of the two systems should have no correlation. (**C**) The belief-spatial location discriminability correlation is positive when spatial location is believed to be relevant (histogram and magenta cumulative distribution curve, p=4×10^−6^), but not when it is believed to be irrelevant (cyan cumulative distribution curve, p>0.05). The two distributions are significantly different (Wilcoxon rank sum test, p=0.014). (**D**) Similarly, belief-spatial frequency discriminability is significantly positive when spatial frequency is believed to be relevant (p=0.0015) but not when it is believed to be irrelevant (p>0.05). The two distributions are significantly different (Wilcoxon rank sum test, p=0.03).

Armed with a neuronal estimate of task-belief and the change in each visual feature on each trial, we looked for interactions between them. Specifically, we tested the hypothesis that strong task-beliefs are associated with stronger encoding of the believed-relevant, but not the believed-irrelevant feature. For each task-belief and stimulus condition, we therefore measured the correlation between belief strength measured from area 7a and perceptual discriminability measured in V1 (Figure 4B). Despite the fact that the resulting correlation is based on few trials and only a few dozen neurons across two very weakly connected areas (Markov *et al*., 2014), we found evidence supporting our hypothesis. There is a positive correlation between belief and the encoding of each feature when it is believed to be relevant, but not when that same feature is believed to be irrelevant (Figure 4C-D). Together, our results indicate that belief-based decision making is an integrated system rather than a separable two-stage computation in which a task decision is followed by a corresponding perceptual decision.

### V1 fluctuations affect belief updating

In addition to the trial-by-trial interaction between belief and perception, two pieces of evidence demonstrate that perception on the previous trial affects how task-belief is updated for the upcoming trial. First, trial information beyond reward outcome affects how task-belief will be updated. On average, the monkeys were more likely to switch tasks after they missed rewards on trials with big changes in the stimulus feature believed to be relevant (Figure S4C). This reliance of belief updating on vision is captured by our ideal observer model which optimally updates task-belief based on historical reward, stimulus, and choices (see Methods and Figure S4A). The ideal observer model consistently predicts the animals’ behavior better than an alternative strategy in which every unrewarded trial affects belief independent of visual and choice experience (Figure S4B). Consistent with findings from studies with similar task structure (Purcell and Kiani, 2016a; Sarafyazd and Jazayeri, 2019), our results demonstrate that belief updating is related to the Bayesian posterior probability that the past perceptual decisions were correct, given the evidence contributing to them.

Second, even for sets of ostensibly identical trials (with the same stimulus, choice, and reward), there was a trial-to-trial relationship between fluctuations in the representations of the visual stimuli and belief updating. We used a neuronal population version of the classic single-neuron neurometric curve (Newsome, Britten and Movshon, 1989) to relate fluctuations in the visual cortex to perceptual decision-making. For groups of trials with the same first stimulus, given the perceptual discriminability decoded from V1 population responses to the second stimulus (as in Figure 4A), we calculated the probability of each feature change direction using logistic regression (Figure 5A, similar to the strategy in (Peixoto *et al*., 2021). The population neurometric curve provided a neuronally-derived confidence about the perceptual choice on each trial. That confidence estimate was related to the animals’ task switching behavior (Figure 5B).

**Figure 5.**
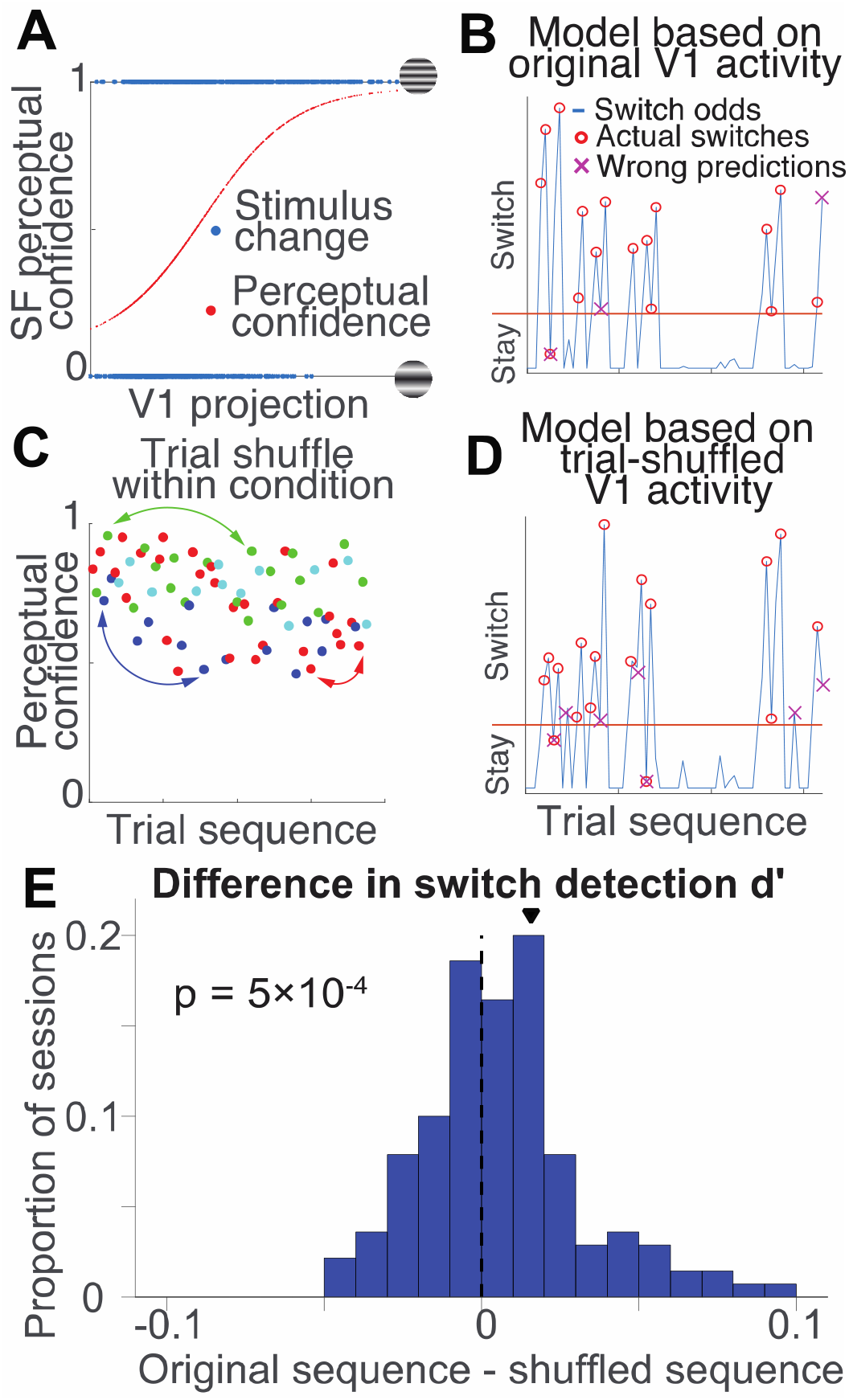
Trial to trial variability in visual cortex affects belief updating. (**A**) Example neurometric curve showing the ability of a decoder to discriminate spatial frequency changes from the population of recorded V1 neurons using logistic regression on the perceptual discriminability of spatial frequency (as in Figure 4A). (**B**) Based on perceptual confidence on each trial (estimated from V1 population activity), a normative model determines whether the subject should switch tasks given the trial history (see STAR Methods). (**C**) Based on the V1 projections to the relevant feature subspace on each trial, we estimate from the neurometric curve, which represents the probability the monkey’s behavioral choice is correct (i.e. perceptual confidence (Hangya, Sanders and Kepecs, 2016)). In the trial-shuffle analysis, we randomly switch the confidence within trials with the same conditions (dots with same color). (**D**) Model predictions after trial-shuffle, conventions as in (b). (**E**) Trial-to-trial variability in V1 is related to belief. The models’ sensitivity indices (*d’*) measures the performances predicting whether the monkey would switch tasks in a session. The histogram shows the differences in the *d’*s of models using the actual and trial-shuffled V1 activity. Across experimental sessions, shuffling V1 activity among trials with identical conditions limited the model’s capability to predict the monkeys task-switching behavior (p=5×10^−4^).

These methods revealed that trial-by-trial fluctuations in visual cortex are related to task switching behavior. A normative model based on the neuronally-derived perceptual confidence on each trial (see STAR Methods) predicts the animals’ behavior better than a comparable model that assumes all trials within the same trial condition (stimuli, choice, outcome) have the same perceptual confidence (Figure S4E). Furthermore, if we shuffle V1 responses among trials with identical trial conditions (Figure 5C), the model’s switch prediction performance suffers significantly (Figure 5D-E). Consistently, unrewarded trials with V1 fluctuations indicating stronger perceptual confidence are more often followed by task switches on the following trial than unrewarded trials with the same trial condition but with V1 fluctuations indicating weaker perceptual confidence (Figure S4D). Together, these results demonstrate that trial-to-trial fluctuations in perception affect belief updating on the subsequent trial, even though these fluctuations provide no benefit for estimating the relevant feature.

## Discussion

Perceptual decision-making has long been studied as an independent cognitive process since it typically occurs on a much faster time scale than the dynamics of higher-level cognitive processes like belief, arousal, and learning. However, our findings demonstrate that there is no such thing as a standalone perceptual decision-making process: every aspect of perceptual decision-making is profoundly integrated with the dynamic task-belief states that dominate natural behavior. We found the relationship between task-belief and perception to be bidirectional.

First, we demonstrated that fluctuating task-beliefs affect perceptual performance. Using a combination of multi-neuron, multi-area physiology, complex but controlled behavior, and hypothesis-driven dimensionality reduction, we demonstrated that perception and task-belief are intimately intertwined such that weak task-beliefs are associated with poor perception of task-relevant information. This suggests that instead of a homogenous mechanism such as arousal, task-belief strength is selectively associated with the processing of believed-relevant information.

The specific cognitive origin of such flexible task-belief-specific modulation remains to be investigated. Potential neuronal mechanisms may include a limit in working memory capacity (Cowan, 2016; Bouchacourt and Buschman, 2019), a reinforcement learning mechanism in perceptual decision-making (Lak *et al*., 2020a; Mendonça *et al*., 2020; Lak *et al*., 2020b), a tradeoff between stable encoding and cognitive flexibility (Driscoll et al., 2017), a neuronal mechanism that limits reinforcement learning to the relevant feature (Niv et al., 2015), flexible geometry of sensory representations (Bernardi et al., 2020; Okazawa et al., 2021), or a feedback-induced change in the structure of correlated variability (Bondy, Haefner and Cumming, 2018). In future work, it will be interesting to determine whether fluctuations in the dynamic weighting of different sources of information (Gu, Angelaki and Deangelis, 2008; Raposo et al., 2012) or other types of beliefs (e.g. those reviewed in Ma and Jazayeri, 2014) interact with perceptual decision-making in similar ways as task beliefs.

Second, we demonstrated that fluctuations in perceptual decision-related neuronal activity affect belief updating. This aspect brings new insights to the current knowledge about cognitive flexibility from studies using unambiguous stimuli, and therefore has very little perceptual involvement (Ravizza and Carter, 2008; Botvinick and Braver, 2015; Bartolo and Averbeck, 2020). Using perceptually challenging tasks is not only more realistic, but also offers powerful tools to investigate, manipulate, and quantitatively understand the specific neuronal process governing perception and belief updating. Our study demonstrates that incorporating these more natural behavioral features into well-controlled laboratory studies leads to important new insights.

In this regard, our findings are merely a starting point of a wide range of potential scientific explorations that will shed light on cognitive flexibility, belief-based decision-making, and learning. For instance, uncertain inference of the task environment would require nonlinear adaptive evidence accumulation (Murphy *et al*., 2021), which could constrain models of individual learning policies (Ashwood *et al*., 2020). These approaches could help uncover the mechanisms underlying the loss of cognitive flexibility in patients with neurodegenerative disorders (Owen *et al*., 1993; Albert, 1996; Gorno-Tempini *et al*., 2004).

The inextricable link between dynamic task-beliefs and perceptual decision-making highlights the importance of studying complex behavior as a whole (Juavinett, Erlich and Churchland, 2018), rather than as stacked building blocks of simpler functions. Complex behaviors likely involve a wide variety of cognitive states other than task-belief, consistent with extensive reports of dynamic cognitive influences on decision-making, including reward expectation (Cicmil *et al*., 2015; Starkweather *et al*., 2017), biases (Akaishi *et al*., 2014; Talluri *et al*., 2018; Mochol, Kiani and Moreno-Bote, 2021), working memory (Panichello and Buschman, 2021), and mood (Yuen and Lee, 2003). Our work therefore showcases a potentially widely applicable approach to study these dynamic, internal cognitive states, using carefully designed cognitive tasks, and by investigating the interaction between these internal states and decision-making on single trials using neuronal population activity. This approach can potentially bring important insights into the fundamental neuronal mechanisms underlying flexible behavior.

Our results also open up exciting avenues for translational therapies that address deficits in flexible decision-making associated with neuropsychiatric disorders. For instance, our results imply that cognitive flexibility is mediated by interactions between neural populations responsible for perception and task-belief. As such, therapies that affect communication between brain areas (e.g. by affecting neurotransmitters like dopamine (Botvinick and Braver, 2015; Mueller *et al*., 2017) have the potential to improve cognitive flexibility in health and disease. Indeed, stimulants that affect the dopamine system like methylphenidate or amphetamines can change focus and flexibility (Bagot and Kaminer, 2014; Mueller *et al*., 2017). Going forward, studying the highly integrated belief-based decision-making system will open up doors to potential treatments of conditions that affect cognitive flexibility and even solutions for healthy individuals to become better decision-makers in volatile environments.

## Acknowledgments

We are grateful to Karen McKracken for providing technical assistance and to Josh Gold, Lori Holt, Douglas Ruff, Amy Ni, Ramanujan Srinath, and Morteza Sarafyazd for comments on an earlier version of this manuscript. This work was supported by the Simons Foundation (Simons Collaboration on the Global Brain award 542961SPI to MRC), the National Eye Institute of the National Institutes of Health (awards R01EY022930 and R01NS121913 to MRC), and the McKnight Foundation (McKnight Scholar award to MRC).

## Author contributions

C.X., and M.R.C. designed research; C.X., and L.E.K. performed research; C.X. performed data analyses and computational modelling; M.R.C. supervised the findings of this work; C.X., and M.R.C. wrote the paper.

## STAR Methods

### Resource Availability

### Key resources table

| REAGENT or RESOURCE | SOURCE | IDENTIFIER |
| --- | --- | --- |
| Deposited data |  |  |
| Original data | This paper | https://doi.org/10.6084/m9.figshare.19591012.v1 |
| Experimental models: Organisms/strains |  |  |
| Rhesus macaque (*Macaca mulatta*) | University of Pittsburgh | N/A |
| Software and algorithms |  |  |
| Original analyses and modelling code | This paper | https://doi.org/10.6084/m9.figshare.19591012.v1 |
| Psychophysics Toolbox | Brainard, 1997 | http://psychtoolbox.org/ |
| MATLAB | The MathWorks. | https://www.mathworks.com/products/matlab.html |
| PsychRNN | Ehrlich *et al.*, 2021 | https://psychrnn.readthedocs.io/en/latest/ |
| Tensorflow | Abadi *et al.*, 2016 | https://www.tensorflow.org |

### Lead contact

Further information and requests for resources should be directed to and will be fulfilled by Marlene Cohen, Department of Neuroscience and Center for Neural Basis of Cognition, University of Pittsburgh, Pittsburgh, PA, 15260, USA.

### Materials availability

This study did not generate new unique reagent.

### Data and code availability

Related data, along with original code, have been deposited at figshare and are publicly available as of the date of publication. Accession numbers are listed in the key resources table.

Any additional information required to reanalyze the data is available from the lead contact upon reasonable request.

#### Experimental subjects

The subjects in our study were two adult male rhesus monkeys (*Macaca mulatta*, monkey F weighed 12 kilograms, monkey G weighed 9 kilograms), who were both experimentally naïve prior to the current experiments. All animal procedures were approved by the Institutional Animal Care and Use Committees of the University of Pittsburgh and Carnegie Mellon University. After we implanted each animal with a titanium head post, they were trained to perform two-interval, two-feature discrimination with stochastic rule switching (Figure 1A-B) (monkey F was trained for 11 months, monkey G for 9 months). We made sure the animals understood the essential requirements of the task based on their behavior (Figure 1C-D), before implanting each animal with 6×8 microelectrode arrays (Blackrock Microsystems) in both parietal cortical area 7a and visual cortical area V1. Each array was connected to a percutaneous connector that allowed daily electrophysiological recordings. The distance between adjacent electrodes was 400 μm, and each electrode was 1 mm long. We implanted 7a arrays on the crown of gyrus between intraparietal sulcus and superior temporal sulcus at approximately 11mm lateral to the midline; and V1 arrays posterior to the lunate sulcus at approximately -13 mm to the intra-aural line and 7 mm lateral to the midline (Figure 1C).

#### Behavioral task

To study perceptual decision making under evolving task-beliefs in dynamic environment, we trained the animals to perform a two-interval, two-feature discrimination task with stochastic task switching. A trial began when the subjects fixated their gaze on a central dot on the screen and they were required to maintain fixation as long as the dot remained on the screen, or the trial would be aborted and unrewarded. A Gabor stimulus was then displayed for 200 ms. After a random delay (300ms to 500ms), a second Gabor stimulus was displayed for 200 ms with a slightly different spatial location (shifted left or right) and a slightly different spatial frequency (higher or lower), with independently randomized change amounts in the two features. The ranges of change amounts are titrated at the beginning of each session so that the overall perceptual performances of the spatial location task and spatial location task are both approximately 75%. Following a subsequent delay of 150ms, the fixation dot disappeared, and the animals were required to make a saccade to one out of four peripheral targets to indicate both the inferred relevant feature and the direction of change in that feature. The two cyan targets correspond to the increase and decrease of spatial frequency when it was believed to be the relevant feature, while the two magenta targets correspond to the left-shift and right-shift of spatial location. The location of the array of targets varied across experimental sessions but remained the same within each session. The monkeys were rewarded only if they correctly reported the direction of change in the relevant feature. The visual stimuli throughout a trial contain no information about the behavioral relevance of features. The relevant feature switches on a randomly chosen 2.5% of trials. The monkeys therefore needed to infer the relevant feature based on their choice and reward history.

#### Electrophysiological recording

All visual stimuli were displayed on a linearized CRT monitor (1,024 × 768 pixels, 120-Hz refresh rate) placed 57 cm from the animal. We monitored eye position using an infrared eye tracker (Eyelink 1000, SR Research) and used custom software (written in Matlab using the Psychophysics Toolbox(Brainard, 1997) to present stimuli and monitor behavior. We recorded eye position and pupil diameter (1,000 samples per s), neuronal responses (30,000 samples per s) and the signal from a photodiode to align neuronal responses to stimulus presentation times (30,000 samples per s) using hardware from Ripple. We recorded neuronal activity from Utah arrays during daily experimental sessions for several months in each animal (90 sessions from monkey F and 62 sessions from monkey G). We set the threshold for each channel at three times the standard deviation and used threshold crossings as the activity on that unit. We positioned the stimuli to maximize the overlap between potential stimulus locations and the joint receptive fields of V1 units, as determined using separate data collected while the monkeys fixated and Gabor stimuli were flashed across a range of retinal positions. Less than 5% of the recorded 7a units had clear receptive fields that we could measure. Those that did had contralateral receptive fields ∼30 degrees in the periphery, which is far away from the Gabor stimuli and the visual targets. We confirmed that during the experiment, the recorded 7a units did not respond selectively to different stimuli (Figure S1B, upper panel).

We included experimental sessions for analyses of neuronal data if they contained at least 480 completed trials (where monkeys successfully maintained fixation until they indicated their choice). We analyzed the activity of area 7a units during the first 300ms after the offset of the first stimulus, when there is no Gabor stimulus on the screen; and the activity of area V1 units during stimulus display periods, shifted with 34 ms visual latency. Units from area 7a were included if their activity during the average delay period was at least 5 sp/s. Units from V1 were included in the analyses if their average stimulus response was 1) at least 25% larger than baseline activity, measured 100ms before stimulus onset, and 2) larger than 5 sp/s. These procedures resulted in 77 sessions from Monkey F and 58 sessions from Monkey G. The median unit counts are 47 in area 7a, 30 in V1. The median trial number in a session is 1086.

#### Recurrent Neural Network

We trained recurrent neural networks to perform the two-interval, two-feature discrimination task with stochastic task switching. Similar to the monkeys, the network model was trained to infer the relevant feature and make the correct perceptual discrimination based on the implicit task rule. The training and testing of the neural network were implemented with custom code based on the open source package PsychRNN (Ehrlich *et al*., 2021) and Tensorflow (Abadi *et al*., 2016). The model consists of 100 units with all-to-all recurrent connections, without any a priori constraints on the connection weights. All results are qualitatively consistent with network sizes of 50 or 200 units.

To produce a choice on each iteration, the recurrent neural network receives input from four channels (Figure 2D): the spatial location change, the spatial frequency change, the recent choice history of the model, and the reward feedback associated with those choices. The recurrent neural network is connected to four output units, which correspond to the four saccade targets presented to the monkey. We take the output unit with the highest activity during choice period as the behavioral choice of the model. Gaussian noises were added to the stimulus changes so that the overall perceptual performances of the model would match those of the monkey (Figure 2E). We trained the model using custom code modified from PsychRNN that allowed the model’s previous choice output to be part of the trial history input for the following trials. The past trial history information was fed into the network in chronological order: each trial starts with the stimulus changes followed by (in order) the choice the model made on the previous iteration, the reward feedback, and finally an intertrial interval before processing information from the next trial. For the model results shown in Figure 2 and Figure S3, the neural network considered a trial history of 7 trials before the current trial, which is longer than most of the monkeys’ task-switch delays. The network behaviors shown in Figure 2 and Supplementary Figure S3 are qualitatively similar if the history input contained 4 trials or 10 trials before the current trial.

The loss function is defined as the squared error between the network output and the target output during the choice period. The target output is 1 on the output unit of the correct choice, and 0 for the other three output units. We initialized the weight matrix with random connections and used a learning rate of 0.001 during training.

#### Population analyses

To obtain a continuous neuronal measure of the animals’ belief state, we analyzed the activity of the population of 7a neurons during the delay period in a high dimensional space in which the activity of each unit was one dimension. We used linear discriminant analysis to identify the best hyperplane to discriminate between 7a population activity on trials where monkeys chose spatial location targets from trials when they chose spatial frequency targets. We defined the belief strength on each trial as the Euclidean distance from the 7a population response to the discriminant hyperplane. Similarly, we obtained a continuous neuronal measure of the discriminability of stimulus change using V1 activity.

#### Statistical tests

All p-values reported in this study are from Wilcoxon signed rank test unless otherwise specified.

#### Normative behavioral model

We use a normative model to characterize belief updating of an ideal observer, given the trial history and the perceptual ability of the monkey (Sarafyazd and Jazayeri, 2019; Purcell and Kiani, 2016a; Glaze *et al*., 2018). Based on the monkeys’ psychometric curve in an experiment session and the change amount of the chosen feature in each trial, we estimated the trial-by-trial probability that their perceptual choice was incorrect. For a non-reward trialed, the odds of likelihoods that the actual task is different from the monkeys’ subjective belief is given by

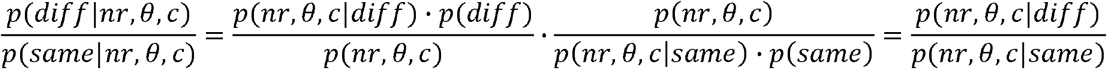

where *θ* and *c* refer to the stimulus change amount and perceptual choice in the feature the monkeys believed to be relevant; and *nr* refers to a non-reward trial outcome. We assumed that overall, the monkeys experienced an equal number of trials where the feature was the same or different from their current belief (i.e., *p* (*diff*) *= p* (*same*)). The monkeys were never rewarded on trials when their subjective task-belief was different from the actual task rule, so *p* (*nr,θ,c*|*diff) =* 1. Meanwhile when subjective task-belief is consistent with the actual rule, the probability of perceptual error can be simply derived from the psychometric function associated with that choice:

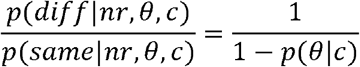

where *p* (*θ*|*c)* is the psychometric function associated with the perceptual choice *c* (Figure 1C). In Figure 4, the psychometric function is replaced with the neurometric function (Figure 4A, also see *Population analyses* section).

For *n* consecutive non-reward trials, the likelihood ratio grows larger as perceptual evidence for a task switch grows as

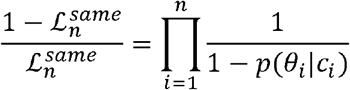

where *ℒ*_*n*_ is the likelihood that the task has not changed after *n* consecutive non-rewarded trials (examples in Figure S4A left panel). Aside from perceptual evidence, the observer presumably also has prior knowledge about the volatility of the task environment. After *n* consecutive non-reward trials, the prior probability that the task stays the same with the last rewarded trial is

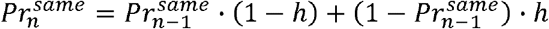

where *h* represents the hazard rate of task change at each trial (for an ideal observer, *h = 0*.*025*, see *behavioral task* section), with

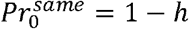

Examples of 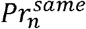 under different environment volatility are shown in Figure S4A middle panel. Taking both perceptual evidence and prior knowledge of environment volatility into account, the model shows that an ideal observer should switch tasks if the posterior probability is higher for actual task rule being different from the subjective task-belief than when they are the same (Figure S4A right panel), i.e.:

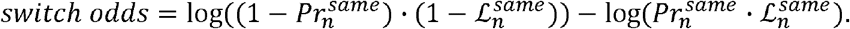

*when positive, switch tssk*

