## Supplementary figures and images for "Dynamic task-belief is an integral part of decision-making"

### Figure S1

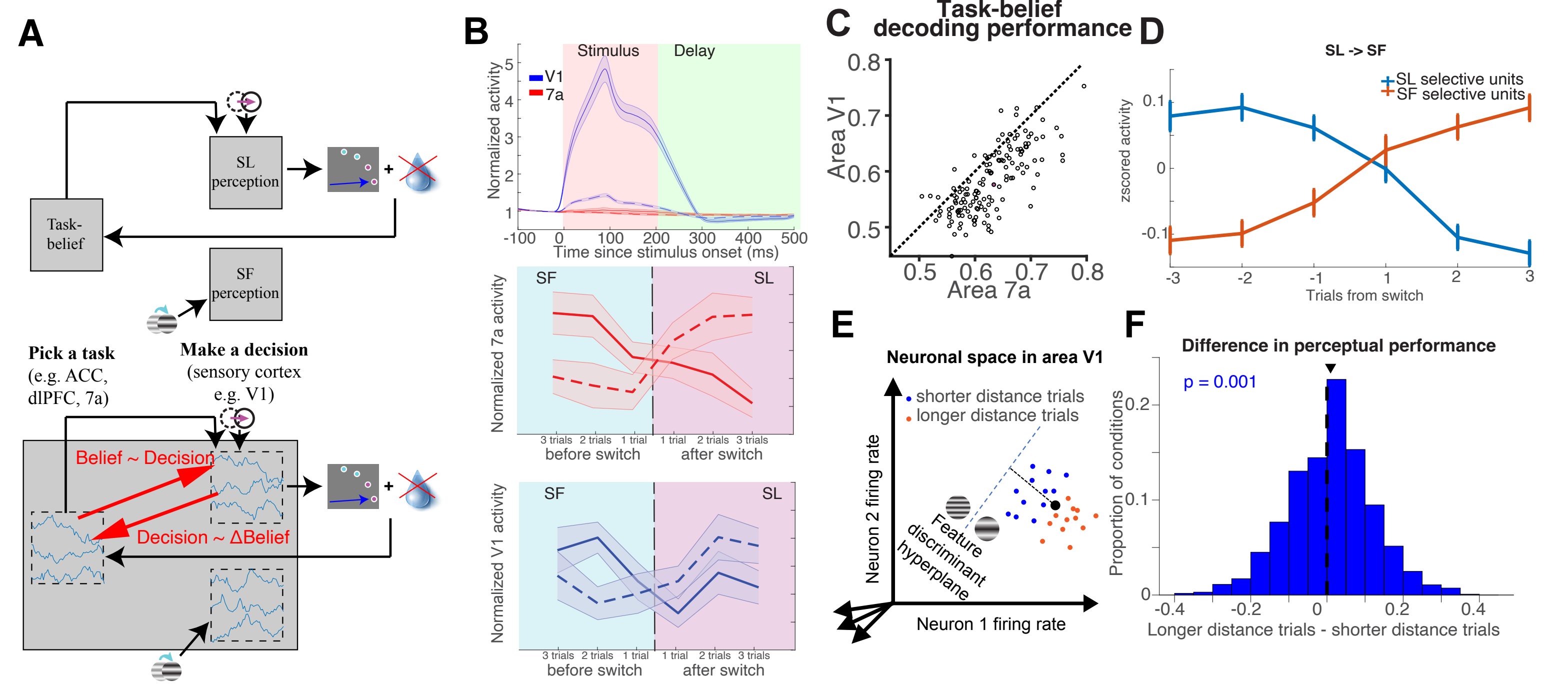

### Figure S2

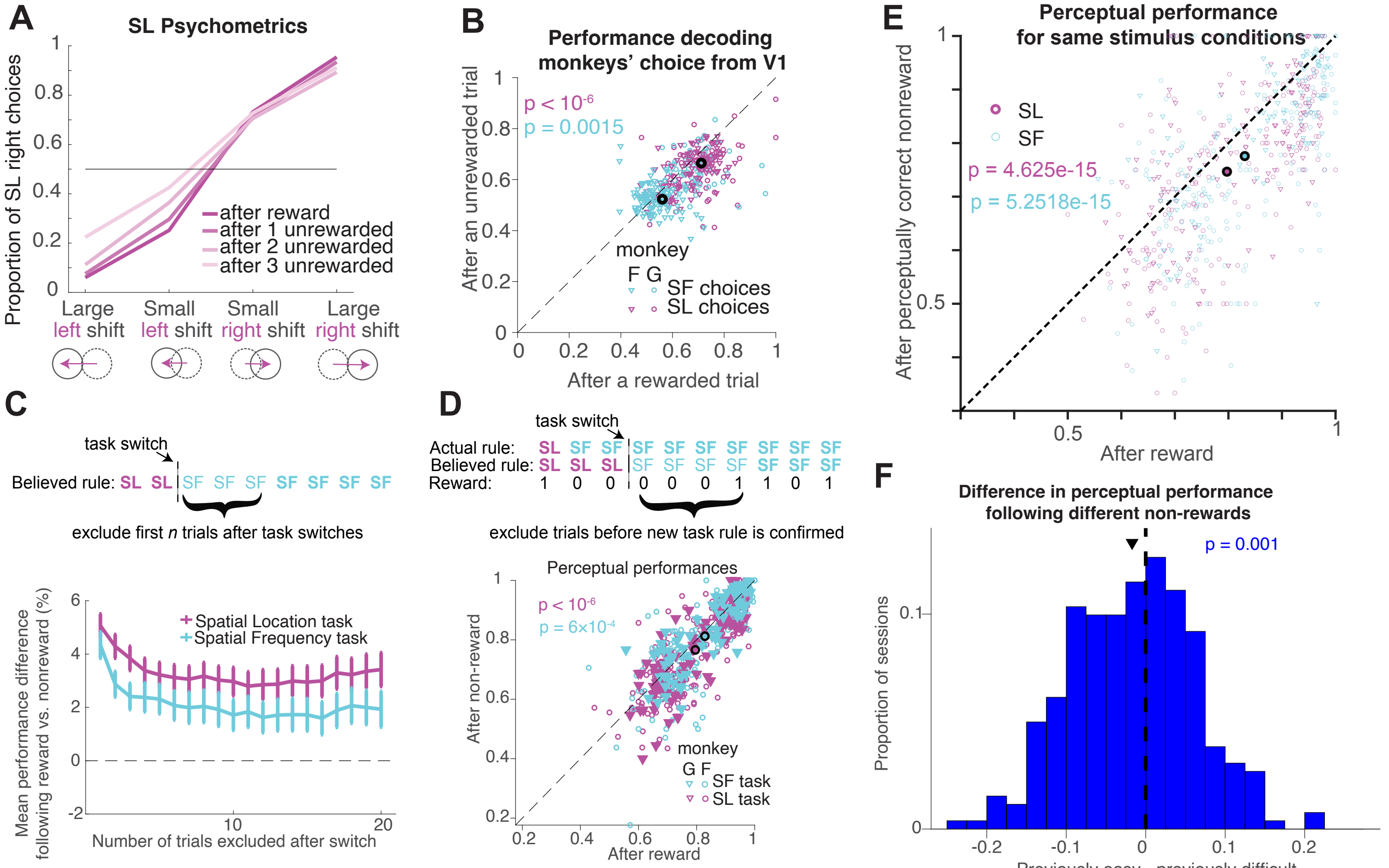

### Figure S3

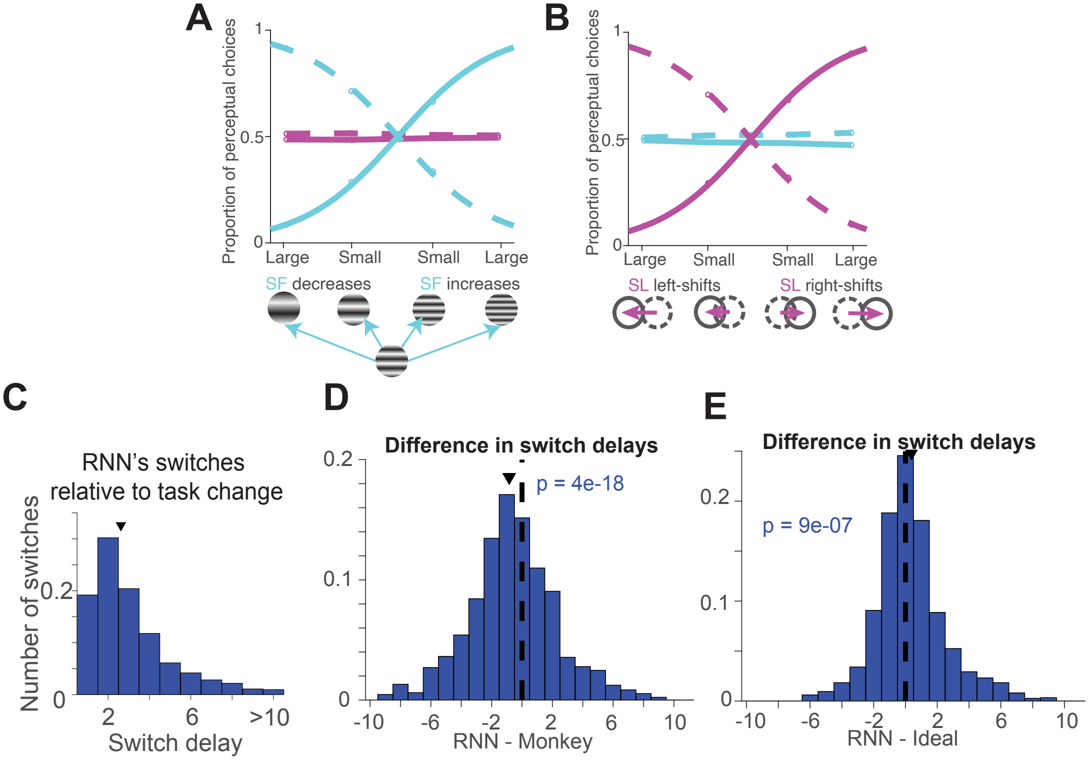

### Figure S4

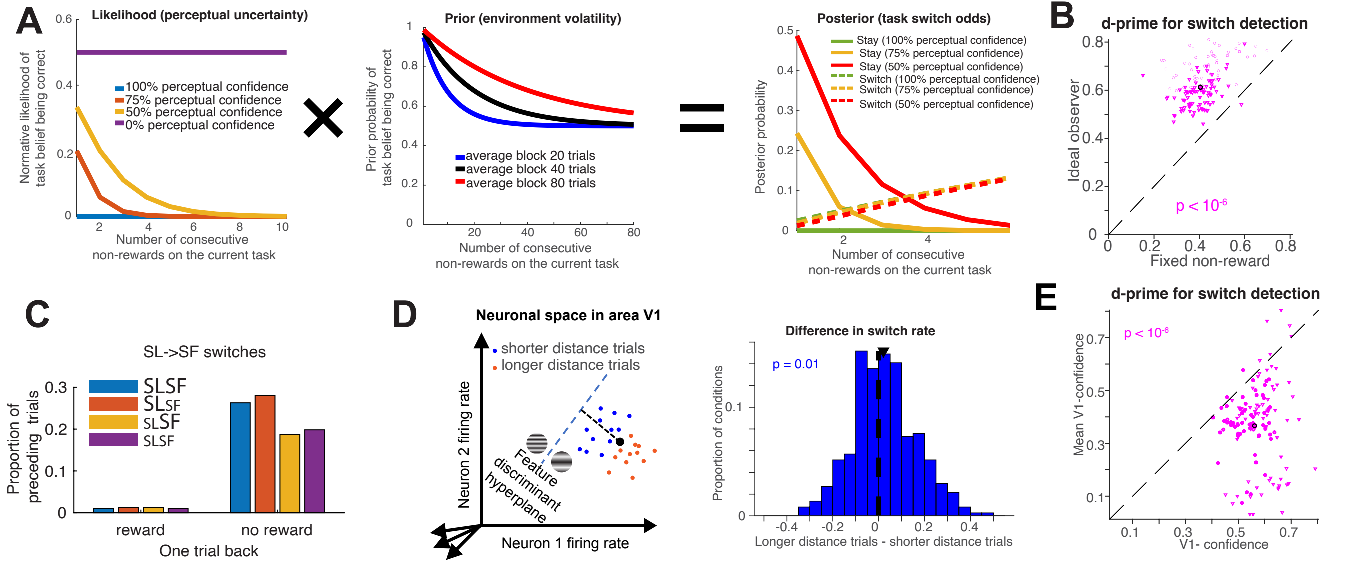
